# A strain-specific metabolic role for the UDP-glucose 4-epimerase Uge3 in *Aspergillus fumigatus* virulence

**DOI:** 10.64898/2026.06.17.732805

**Authors:** Nicole E. Kordana, Angus Johnson, Charles Puerner, Jane T. Jones, Caitlin H. Kowalski, Katherine G. Quinn, Ko-wei Liu, François Le Mauff, Robert A. Cramer

**Affiliations:** Geisel School of Medicine, Microbiology and Immunology Department, Dartmouth College, Hanover NH, USA; Infectious Disease and Immunity in Global Health Program, Research Institute of the McGill University Health Centre, Montreal, Quebec, Canada; GlycoNet Integrated Services, Microbial Glycomic Node, Montreal, Quebec, Canada

**Author notes:** Corresponding Author: Robert A. Cramer, Ph.D. AJ and CP equally contributed to this study.

## Abstract

Expression of a fungal-specific sub-telomeric gene, *hrmA*, in *Aspergillus fumigatus* is important for a colony biofilm morphology termed H-MORPH, increased hypoxic fitness, and virulence in a murine model of invasive pulmonary aspergillosis (IPA). How expression of *hrmA* contributes to virulence and worse disease progression is ill-defined. Increased *hrmA* expression results in reduced attachment of the extracellular matrix (ECM) to the fungal cell wall resulting in decreased strain adherence. Fungal strains that are less adherent *in vitro* are typically less virulent as the ECM heteropolysaccharide galactosaminogalactan (GAG) aids in adhesion to host cells and confers protection from host responses. Here we report that the UDP-glucose 4-epimerase encoding gene required for GAG biosynthesis, *uge3*, is necessary for full virulence of the H-MORPH strain, *hrmA*^REV^ (AF293::*hrmA*^D304G^). In contrast, loss of *uge3* in the reference strain AF293 did not significantly impact virulence in the tested IPA murine model. Phenotypic, transcriptomic, and metabolic analyses of *uge3* loss in the respective strain backgrounds revealed a key role for Uge3 in central carbon metabolism in a strain specific context that promotes disease progression. These results complement the known role of Uge3 in GAG biosynthesis and highlight strain specific metabolic differences in pathogenic *A. fumigatus* strains.

**IMPORTANCE:** *Aspergillus fumigatus* forms adherent biofilms that contribute to its ability to persist and cause disease. However, significant strain diversity exists with regard to the morphology of *A. fumigatus* biofilms. A distinct colony morphotype associated with increased disease progression and low oxygen fitness, termed H-MORPH, was recently described. An additional defining feature of the H-MORPH biofilm morphotype is reduced *in vitro* adherence to surfaces. While reduced fungal strain adherence is most commonly associated with reductions in virulence, H-MORPH strains exhibit increased virulence relative to the well-studied N-MORPH reference strain AF293. Here we discover that the UDP-glucose 4-epimerase, Uge3, plays an important role in H-MORPH central carbon metabolism complementary to its role in production of the extracellular matrix polysaccharide galactosaminogalactan (GAG). In H-MORPH strains, this metabolic role for Uge3 becomes central to virulence. These data highlight *A. fumigatus* strain specific mechanisms of fungal carbon metabolism related to biofilm matrix production and fungal virulence.

## INTRODUCTION

High mortality mycoses caused by the mould *Aspergillus fumigatus* pose an increasing threat to human health in the face of limited treatment options, rising antifungal resistance, and novel susceptible patient populations (1–4). Thus, understanding fungal physiology during infection is important to identify fungal targets for therapeutic development. A key feature defining *A. fumigatus* infection microenvironments is oxygen limitation (5). Subsequently, adaptation to low oxygen environments is essential for *A. fumigatus* virulence and strains exhibiting increased low oxygen fitness correlate with worse disease outcomes in murine models of invasive pulmonary aspergillosis (IPA) (6, 7). As oxygen is essential for cellular processes governing fungal growth such as ATP-production and sterol biosynthesis, low-oxygen adaptation through a hypoxia response necessitates metabolic reprogramming that impacts fungal physiology (8, 9). In *A. fumigatus*, acute exposure and chronic adaptation to low oxygen environments also alters the cell wall, highlighted by increased beta-glucan exposure (10, 11). *In vitro* and *in vivo*, hypoxia response-driven increases in fungal beta-glucans are associated with regions of reduced oxygen availability and increased host inflammatory responses that are at least partially dependent on the host C-type lectin receptor, Dectin-1 (*Clec7a*) (10–12). However, it is not known if increased host recognition of exposed fungal beta-glucans by Dectin-1 contributes to increased virulence exhibited by low oxygen-fit *A. fumigatus* strains.

Another key feature of low oxygen adapted *A. fumigatus* strains is decreased *in vitro* adherence (11, 13). Strain adherence is attributed to biosynthesis of the major component of the extracellular matrix (ECM), galactosaminogalactan (GAG), which is an established virulence factor of *A. fumigatus* (14–16). GAG is a heteropolymer comprised of galactose (Gal) and partially deacetylated N-acetylgalactosamine (GalNAc); however, other polysaccharide components of the ECM include α-glucans (glucose, Glc) and galactomannan (galactose, Gal; mannose, Man) (17–21). The five genes necessary for GAG production were identified from transcriptomic analyses of transcription factor mutants with impaired biofilm adherence (14). Our current understanding of GAG biosynthesis begins intracellularly where the bifunctional enzymatic activity of the UDP-glucose 4-epimerase, Uge3, can epimerize UDP-Glc to UDP-Gal and UDP-GlcNAc to UDP-GalNAc (14, 19). These nucleotide sugars are thought to be polymerized and exported from the fungal cell via the large transmembrane glycosyltransferase, Gtb3. Once outside of the cell, the three remaining enzymatic genes in the cluster, Ega3, Sph3, and Agd3, mediate polysaccharide processing via specific sugar cleavage events and partial deacetylation to render a cationic polysaccharide capable of adhering to the cell wall and inert surfaces via charge-charge interactions (22–24).

GAG increases adherence to host cells, conceals inflammatory beta-glucans at the fungal cell wall from host detection, and influences the host response through NLRP3 activation, IL-1 production, and resistance to neutrophil extracellular trap formation (14–16, 25). Loss of either *uge3* or *agd3* results in a significant loss of virulence of the AF293 reference strain in murine IPA models, linking GAG production and strain adherence with virulence (14, 22). With a growing body of literature highlighting phenotypic variation across the *A. fumigatus* population, it remains unclear if GAG production varies between strains or how altered production of GAG may contribute to strain specific host responses and ultimately disease (26–29).

In previous work, we identified an allele of the sub-telomeric gene *hrmA* (*hrmA*^D304G^) that is sufficient to increase low oxygen (hypoxic) fitness in the parental strain, AF293 (11). Expression of this allele generates a furrowed colony biofilm morphology termed H-MORPH, that is distinctive from the flat normoxic morphology (N-MORPH) exhibited by the reference strain AF293 (11). H-MORPH is observed in clinical isolates and is highlighted paradoxically by reduced *in vitro* adherence but increased virulence in a murine model of IPA. Despite identifying a fungal gene, *hrmA,* sufficient to induce these phenotypes, it has remained unclear how H-MORPH strains increase virulence in the setting of steroid mediated immune suppression murine models. A hallmark of H-MORPH virulence in this model is increased neutrophil recruitment to the airways, possibly resulting in immunopathogenesis (11). Here we leverage a set of isogenic strains to investigate mechanisms of H-MORPH reduced adherence and increased virulence. Our data suggest that the UDP-glucose 4-epimerase, Uge3, supports H-MORPH virulence through mediating increased hexosamine biosynthetic pathway activity during rapid, proliferative growth, complementing Uge3’s established role in GAG production. We postulate that *A. fumigatus* Uge3 functions similarly to the *Homo sapiens* homolog, GALE, playing a critical role in carbon regulation that we have identified to be important for H-MORPH colony morphology, fungal biomass proliferation, and virulence (19).

## RESULTS

### Host Dectin-1 recognition of fungal cell wall beta-glucans does not explain H-MORPH strain hrmA^REV^ increased virulence

We previously reported the H-MORPH strain, *hrmA*^REV^, exhibits hypoxia response phenotypes in normoxia including a beta-glucan rich cell wall and decreased *in vitro* adherence with reduced hyphal ECM attachment. (11). These phenotypes, taken together with the increase virulence highlighted by a significant increase in neutrophil numbers observed in murine airways challenged with *hrmA*^REV^, led us to hypothesize that exposed beta-glucans drive increased H-MORPH virulence through immunopathogenesis.

To test if the cell wall exposed beta-glucans were driving increased virulence of *hrmA*^REV^, we challenged triamcinolone acetonide immunosuppressed wildtype (C57BL/6J, littermate controls) or *Clec7a*^-/-^ (encoding the Dectin-1 pattern recognition receptor (PRR)) mice with a low dose of conidia (10^5^ /mouse) from either AF293 or *hrmA*^REV^. Dectin-1 is a major PRR of fungal beta-glucan (30–32). If Dectin-1 recognition of fungal beta-glucans is necessary to drive increased virulence of *hrmA*^REV^ we would expect to observe no significant difference in virulence between the strains in *Clec7a*^-/-^mice. However, similar to what we previously observed in an outbred CD-1 mouse genetic background, we observed that *hrmA*^REV^ exhibited increased virulence in both wildtype and *Clec7a*^-/-^ mice (**Fig. 1A**) (11). Upon histological examination of fungal infected lung tissues, we observed that unlike in wildtype C57BL/6J mice, *hrmA*^REV^ was able to invade the host lung tissue in *Clec7a*^-/-^ animals, suggesting Dectin-1 recognition of cell wall beta-glucans in a wildtype mouse is necessary to prevent fungal invasion of *hrmA*^REV^ (**Fig. 1B**). However, histopathological analyses did not present clear differences in inflammation between the different mouse strains challenged with AF293 or *hrmA*^REV^. Consequently, it is unlikely that increased Dectin-1 mediated host cell recognition of *hrmA*^REV^ is responsible for its increased virulence relative to AF293.

**Figure 1.**
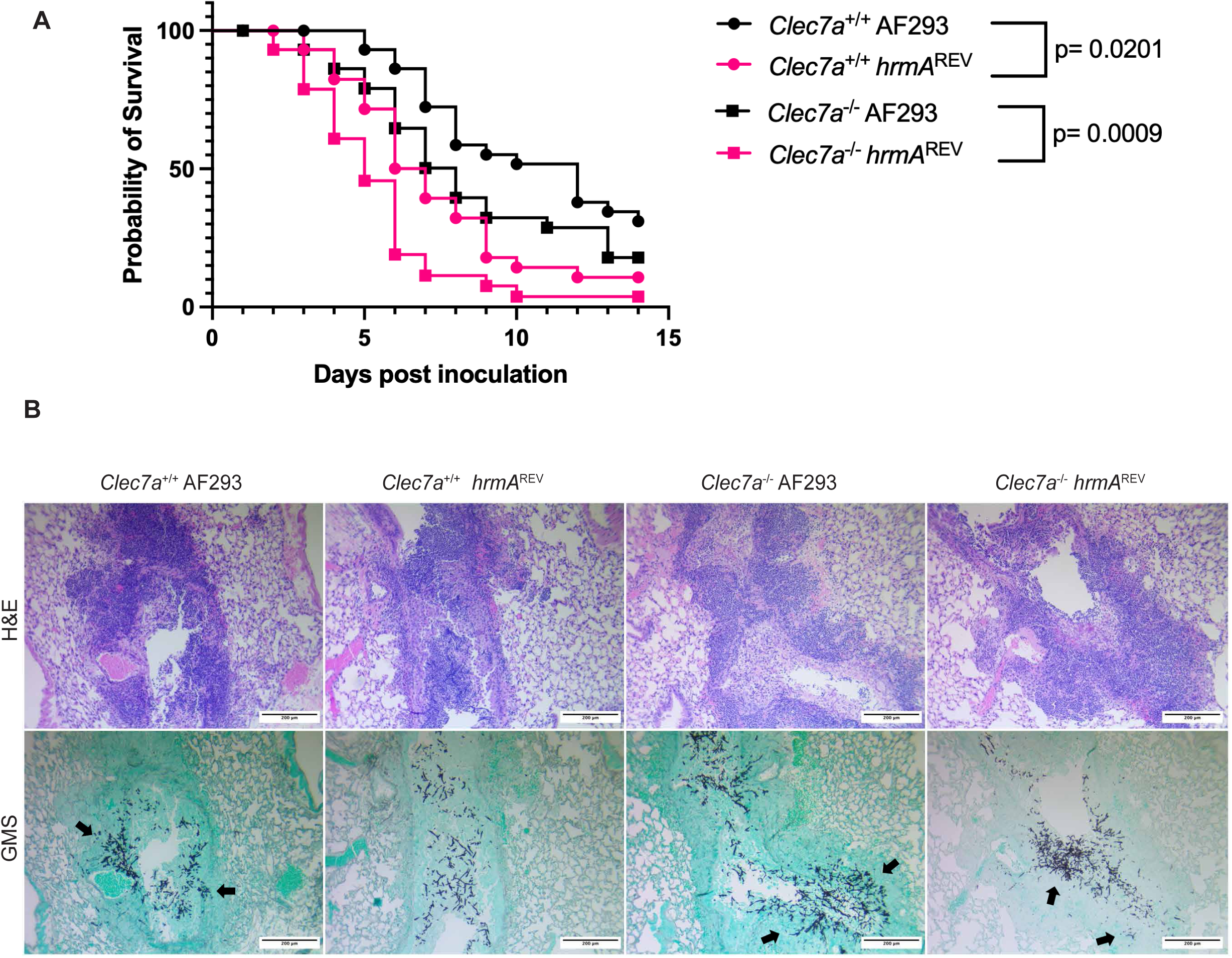
Host recognition of fungal beta-glucans does not explain increased virulence of the H-MORPH strain *hrmA*^REV^. *hrmA*^REV^ is significantly more virulent than the N-MORPH strain, AF293, in both wildtype and *Clec7a*^-/-^ mice. **A)** Wildtype or *Clec7a*^-/-^ mice (C57B6/J background) 8-12 weeks old, received 40mg/kg Kenalog 24 hours prior to inoculation with 1x10^5^ spores in 40µL of PBS intranasally. Mice were monitored for 14 days post inoculation for mortality. Data represents three independent experiments with each group containing n=10 mice/experiment (n=30 mice/group total). Log-rank (Mantel-Cox) analyses on combined biological replicate data. **B)** Representative histology images (H&E and GMS) of wildtype and *Clec7a*^-/-^ mice that received 10^5^ fungal spores intranasally of either AF293 or *hrmA*^REV^ from 3 days post inoculation. Black arrows indicate fungal invasion into lung tissue. Imaged on Zeiss Axioskop2 Plus utilizing SPOT (v5.2) imaging software scale bars = 200µm.

### hrmA^REV^ produces significantly less GAG with an altered composition compared to AF293

While other host PRRs can detect and respond to fungal beta-glucans (33–37), we next asked if the GAG fraction of the ECM produced by *hrmA*^REV^ contributes to its increased virulence. As it is unknown why the ECM is not well adherent to the *hrmA*^REV^ hyphae, we first aimed to address if the GAG produced by *hrmA*^REV^ is altered in quantity or composition. To address this, we analyzed the secreted polysaccharide fraction of whole cell culture supernatants from AF293 and *hrmA*^REV^ by GC-MS as previously described (38). This approach quantifies all secreted sugars, so in addition to GAG monosaccharides (galactose, Gal; N-acetylgalactosamine, GalNAc), we can detect monosaccharides associated with α-glucans (glucose, Glc) and galactomannan (mannose, Man; galactose, Gal). Additionally, knowing that AF293 and *hrmA*^REV^ exhibit baseline differences in growth and biomass production, fungal biomass was collected and lyophilized to normalize the extracted extracellular matrix to the total amount of fungus in each sample. *hrmA*^REV^ secretes significantly less polysaccharides relative to biomass produced than its parental strain AF293 (**Fig. 2A**). Among these polysaccharides, GAG was strongly affected, with *hrmA*^REV^ having significantly decreased amounts of both Gal and GalNAc sugars compared to AF293 (**Fig. 2A**).

**Figure 2.**
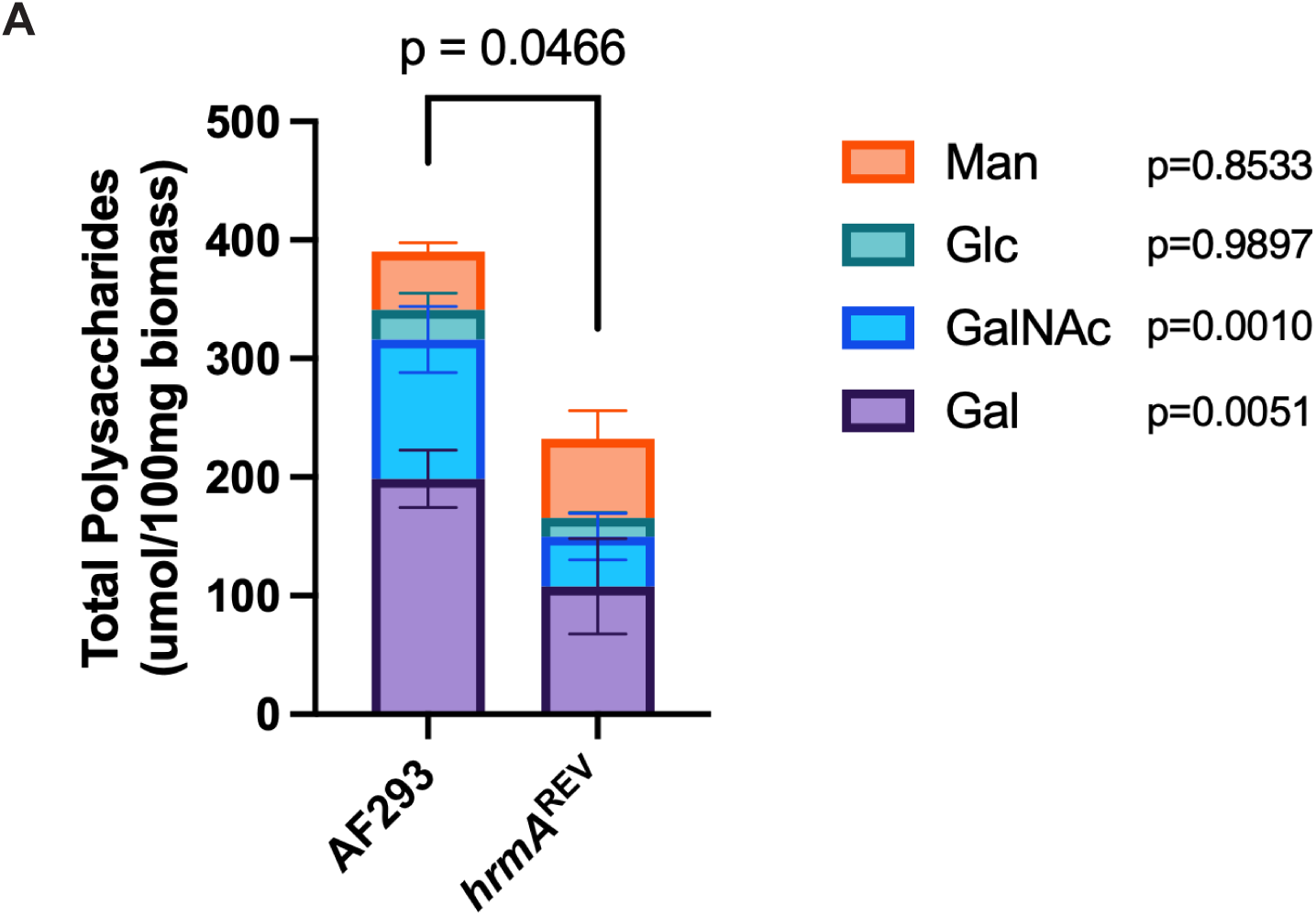
H-MORPH secretes less GAG that is of altered composition. **A)** *hrmA*^REV^ secretes less extracellular sugars than parental AF293. Total extracellular polysaccharides were obtained from whole culture supernatant by dialysis and lyophilization prior to derivatization and quantification via GC-MS. Total extracellular polysaccharides normalized to total fungal biomass produced reveals that *hrmA*^REV^ secretes significantly less polysaccharides compared to parental AF293 (two-tailed Welch’s T-test, p=0.0466). GAG monosaccharides, Gal and GalNAc, are significantly reduced in *hrmA*^REV^ compared to AF293 as determined by ordinary two-way ANOVA with Sidak’s multiple comparisons (Man p=0.8533; Glc p=0.9897; GalNAc p=0.0010, Gal p=0.0051). Data is representative of three independent biological replicates; error bars depict standard deviation of the mean for each quantified monosaccharide.

Neither mannose nor glucose displayed a significant difference between the two strains. The alteration in GAG abundance and composition in combination with previously characterized differences at the cell wall is consistent with the decreased adherence of *hrmA*^REV^, but raises intriguing questions related to strain-specific GAG biosynthesis and function in relation to fungal strain virulence (11). As *hrmA*^REV^ exhibited a reduction of GAG associated secreted polymers, we next sought to address the role of GAG production in H-MORPH virulence.

### Loss of the UDP-glucose 4-epimerase, uge3, decreases characteristic H-MORPH phenotypes and reduces growth

To investigate the role of GAG in *hrmA*^REV^ H-MORPH virulence, we first generated an *hrmA*^REV^Δ*uge3* strain that lacked the ability to produce GAG, along with the reconstituted strain, *hrmA*^REV^Δ*uge3*^recon^. Both the null mutant and reconstituted strain were confirmed via qRT-PCR expression of *uge3* and by crystal violet adherence assays (**Fig. S1A-B**). The AF293Δ*uge3* strain utilized was a generous gift of the Sheppard Lab (14). We observed that *hrmA*^REV^Δ*uge3* had an altered colony biofilm morphology compared to *hrmA*^REV^ and *hrmA*^REV^Δ*uge3*^recon^ (**Fig. 3A**). The alterations to the *hrmA*^REV^Δ*uge3* colony biofilm resulted in decreased characteristics of H-MORPH including reduced furrowing and a reduction in vegetative mycelia (**Fig. 3C-D**) (11). We additionally observed that *hrmA*^REV^Δ*uge3* colony biofilms have a minor growth defect measured by colony diameter compared to *hrmA*^REV^ that is not observed in the submerged liquid culture biofilm model at the time point tested (**Fig. 3B, E**). In contrast, the AF293Δ*uge3* strain did not exhibit a change in colony biofilm morphology or growth (**Fig. 3A-E).** Importantly, *hrmA*^REV^ displayed decreased adherence relative to AF293 as previously reported, and adherence exhibited by all strains is attributed to Uge3 function (**Fig. 3F**). The change in *hrmA*^REV^Δ*uge3* colony morphology, coupled with the slight reduction in colony biofilm growth, suggests an unexpected role for Uge3 in H-MORPH fitness.

**Figure 3.**
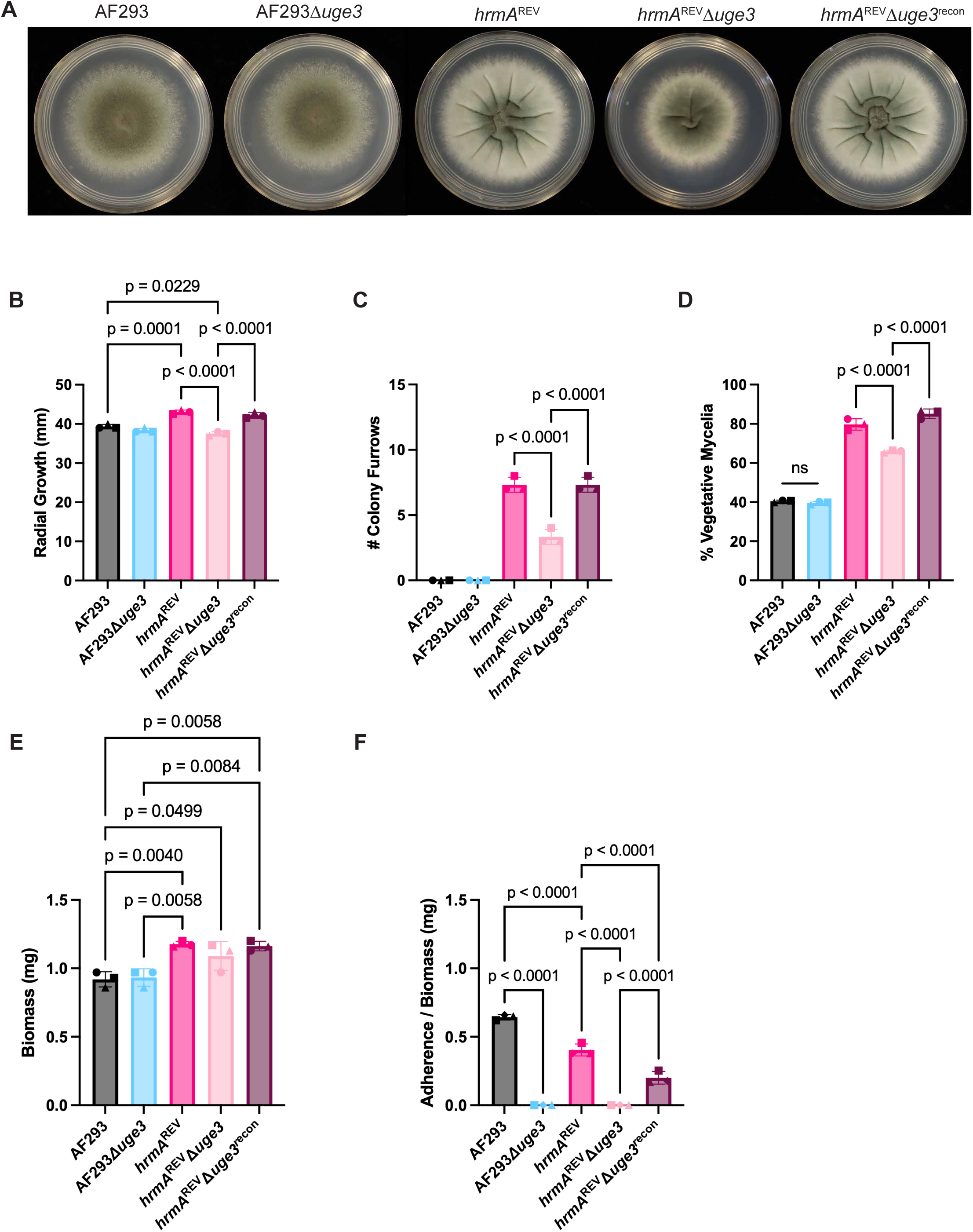
Loss of *uge3* impacts H-MORPH colony morphology. **A-D)** Representative images of 72-hour colony biofilms utilized for quantifying impact to H-MORPH colony morphology via radial growth, number of colony furrows, and percent vegetative mycelia. Data depict three independent biological replicates. **E-F)** Paired biomass and crystal violet adherence data obtained from 24hr submerged biofilms (1x10^5^ spores/mL in LGMM). Data represents three independent biological replicates. **(B, E, F)** Statistical significance assessed via one-way ANOVA with Tukey’s multiple comparisons, **(C-D)** one-way ANOVA with Sidak’s multiple comparisons.

### Loss of uge3 significantly reduces hrmA^REV^ virulence in an immunosuppressed murine model of invasive pulmonary aspergillosis

To evaluate if *uge3* contributes to *hrmA*^REV^ virulence, we next challenged triamcinolone acetonide immunosuppressed CD-1 mice with a low dose (10^5^) of fungal conidia from either AF293, AF293Δ*uge3*, *hrmA*^REV^, *hrmA*^REV^Δ*uge3*, or *hrmA*^REV^Δ*uge3*^recon^. In this murine model of IPA, AF293Δ*uge3* did not display a significant reduction in virulence compared to the parental AF293 (14). In contrast, a significant and striking reduction in virulence was observed for *hrmA*^REV^Δ*uge3* compared to *hrmA*^REV^ and the *hrmA*^REV^Δ*uge3*^recon^ reconstituted strain (**Fig. 4A**). These data strongly suggest *uge3* contributes to virulence in the *hrmA*^REV^ background.

**Figure 4.**
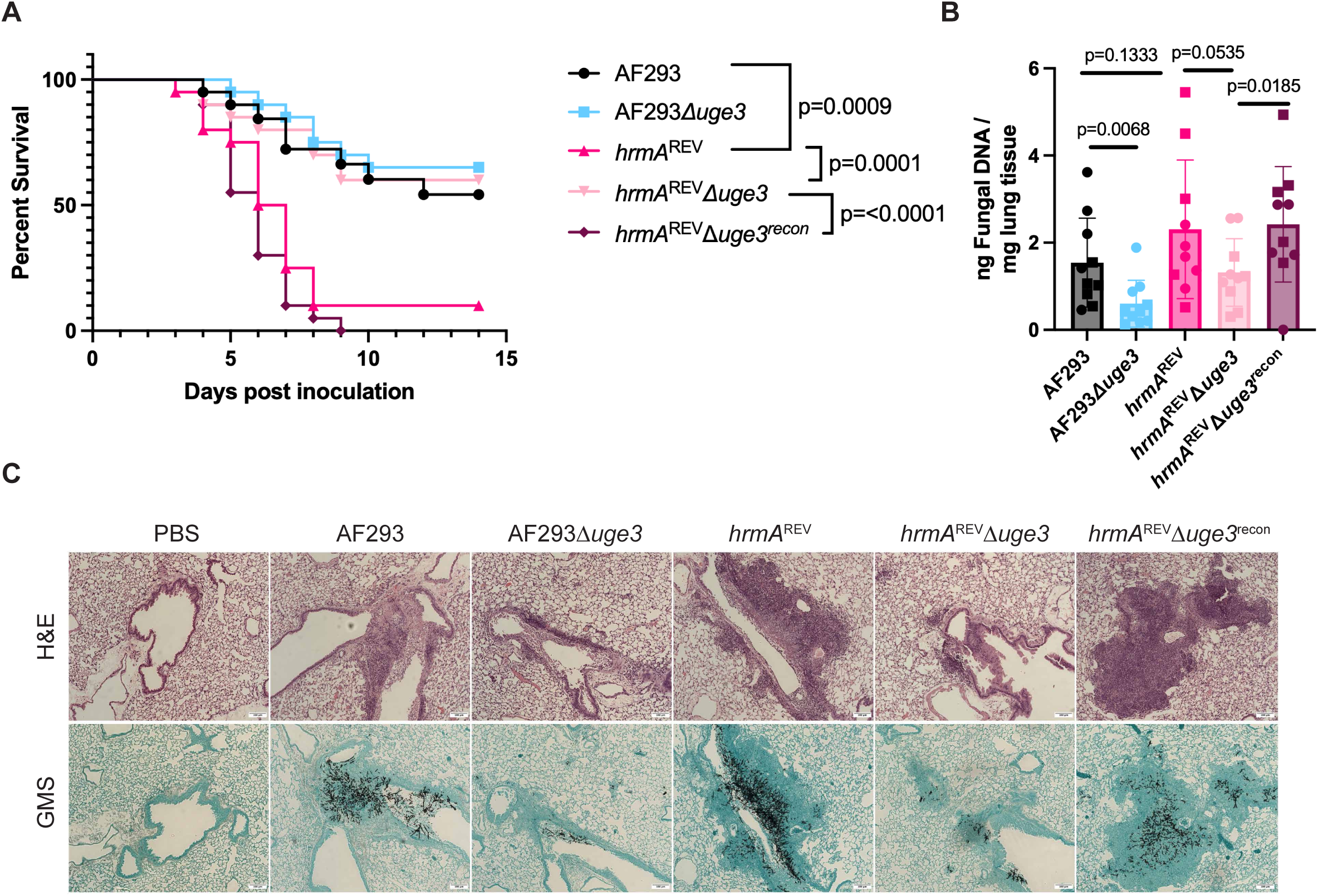
Loss of *uge3* significantly impairs virulence in H-MORPH in an immunosuppressed murine model of IPA. **A)** Kaplan-Meier Survival curve of mice challenged with AF293 or HrmA^REV^ and corresponding GAG-deficient (Δ*uge3*) mutant strains. 6–8wk female CD-1 mice, were immunosuppressed with 40mg/kg triamcinolone acetonide (Bristol-Myers Squibb) 24 hours prior to intranasal inoculation with 1x10^5^ fungal spores in 40µL PBS. Mice were monitored for signs of mortality and morbidity for 14 days. Data represents two independent biological experiments, each with n=10 mice/group. Statistical significance determined utilizing Kaplan-Meier survival analysis, Logrank (Mantel-Cox). **B)** Fungal burden at 3 days post inoculation from corticosteroid immunosuppressed mice that received 1x10^5^ spores in 40µL PBS I.N. Fungal DNA from lungs quantified using qPCR of ITS1 genomic locus, then total fungal DNA extracted normalized to total lyophilized lung tissue weight. Data represents two independent biological experiments, each with n=5 mice/group; Kruskal-Wallis with Dunn’s multiple comparisons. **C)** Histopathology images (10X magnification; H&E and GMS) of corticosteroid immunosuppressed murine lungs 3 days post inoculation with 1x10^5^ fungal spores in 40µL PBS, or 40µL PBS. Images are representative of two independent biological experiments with n=4 mice/group. Imaged on Zeiss Axioskop2 Plus utilizing SPOT (v5.2) imaging software, scale bars = 100µm.

To begin to understand how loss of *uge3* impacts *hrmA*^REV^ virulence, fungal burden and histological examination of lung tissues at 3 days post challenge were assessed. As we previously reported, we did not observe significant differences in fungal burden between AF293 and *hrmA*^REV^ at time points examined (**Fig. 4B**) (11). In alignment with previous reports, AF293Δ*uge3* displayed a significant reduction in fungal burden (**Fig. 4B**), corresponding to a reduction in observed fungal lesions in the lung tissues of challenged mice (**Fig. 4C**) (14). We also observed a significant reduction in fungal burden of mice challenged with the *hrmA*^REV^Δ*uge3* mutant compared to both *hrmA*^REV^ and *hrmA*^REV^Δ*uge3*^recon^ (**Fig. 4B**). Histological examination corroborated this finding, as fewer fungal lesions were observed in the *hrmA*^REV^Δ*uge3* recipient mice compared to mice that received *hrmA*^REV^ or the reconstituted strain. Furthermore, the tissues from mice challenged with *hrmA*^REV^ displayed foci of infection heavily surrounded by neutrophilic inflammation, which was dramatically reduced in *hrmA*^REV^Δ*uge3* and restored in the reconstituted strain (**Fig. 4C**). Taken together these data suggest *uge3* is important for fungal fitness in the lung irrespective of strain background, but essential for virulence in the H-MORPH strain *hrmA*^REV^ in this immunosuppressed murine model of IPA.

### Uge3 supports a pseudohypoxic metabolic state in hrmA^REV^

The finding that *uge3* plays a unique role in mediating both colony morphology and virulence in *hrmA*^REV^ was striking. This led us to ask what is different about *hrmA*^REV^ that loss of *uge3* impacts this strain significantly more than AF293. Homologs of *uge3* are present in both eukaryotes and prokaryotes and play important roles in glucose and galactose metabolism as well as LPS production (39–43). Furthermore, *A. fumigatus uge3* homologs in other *Aspergillus* species also function in the production of nucleotide sugars required for cell wall biosynthesis and GAG production (44–47). UDP-GlcNAc, generated through the hexosamine biosynthetic pathway (HBP), is a substrate for Uge3 to generate GAG (19, 48–50). Given the link between Uge3 and metabolism, we hypothesized *hrmA*^REV^ exhibits an altered metabolic state that relies on Uge3 activity. To test this hypothesis, we conducted RNA-Sequencing (RNA-Seq) on mature submerged biofilms from strains AF293, AF293Δ*uge3*, *hrmA*^REV^, and *hrmA*^REV^Δ*uge3.* Hierarchical clustering of Euclidean distances on VST-normalized counts across samples revealed samples clustered together by strain background with minor variability in clustering by mutant status (**Fig. S2A**). To begin to address how *uge3* may uniquely impact *hrmA*^REV^, we identified significantly differentially expressed genes (DEG) (CPM>10, FDR ≤ p=0.05) between *hrmA*^REV^ vs. AF293 and between *hrmA*^REV^Δ*uge3* vs. AF293Δ*uge3* (**Fig. 5A**). From these analyses we were able to obtain lists of DEGs whose transcripts were significantly increased or decreased due to strain background versus what is uniquely impacted by the loss of *uge3* (**Table S1**). We considered DEGs listed in the center panel of the Venn diagram to be impacted by differences in the strain background as the genes were differentially abundant independent of the null mutant status. DEGs highlight transcripts of genes broadly involved in metabolism to be present in both increased and decreased comparisons (**Table S1**).

**Figure 5.**
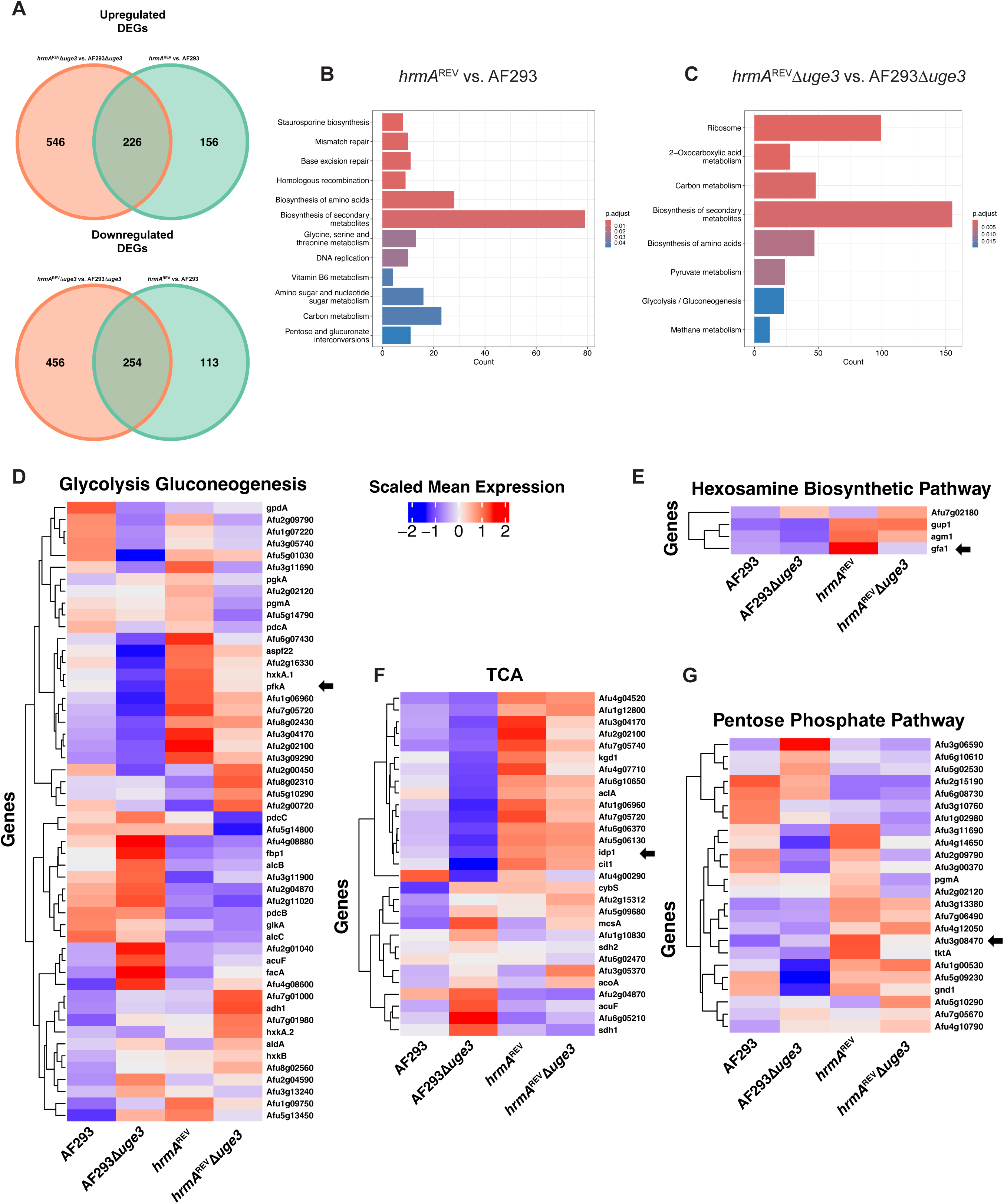
Uge3 supports a pseudohypoxic state in *hrmA*^REV^. **A)** Venn diagrams of upregulated and downregulated DEGs observed in comparisons between *hrmA*^REV^ vs. AF293 and *hrmA*^REV^Δ*uge3* vs. AF293Δ*uge3* (CPM>10, FDR p < 0.05). Gene IDs for these comparisons are listed in **Table S1**. **B-C)** KEGG enrichment analyses of DEGs. **D-G)** Scaled CPM heatmaps of genes (CPM>10, FDR p = <0.05) from glycolysis and gluconeogenesis, TCA, PPP, and HBP. Scaled mean expression key in panel D is applicable to panels D-G. Black arrows indicate gene of the rate limiting enzyme for each pathway.

To better understand changes in metabolism that might be occurring we conducted KEGG pathway enrichment analyses (**Fig. 5B-C**). We observed enrichment in DEGs involved in “carbon metabolism” between *hrmA*^REV^ and AF293 that were more significantly enriched in the comparison between *hrmA*^REV^Δ*uge3* and AF293Δ*uge3* (**Fig. 5B-C**). This suggests that central carbon metabolism differs between *hrmA*^REV^ and AF293 and is exacerbated by the loss of *uge3*. KEGG enrichments for “pentose and glucuronate interconversions” and “amino sugar and nucleotide sugar metabolism” were unique to the comparison between *hrmA*^REV^ and AF293, suggesting the pentose phosphate pathway (PPP) and hexosamine biosynthetic pathway (HBP) may be uniquely important in *hrmA*^REV^. Meanwhile, KEGG enrichments for “pyruvate metabolism” and “glycolysis/gluconeogenesis” were unique to the comparison between *hrmA*^REV^Δ*uge3* and AF293Δ*uge3*, suggesting glycolysis and the citric acid cycle (TCA) may be uniquely impacted by loss of *uge3*.

To further explore these data, we generated heat maps for genes annotated to be involved in glycolysis and gluconeogenesis, TCA, PPP, and HBP (**Fig. 5D-G**). Notably, we found that genes encoding rate limiting enzymes (indicated with black arrows) for glycolysis (*pfkA*; phosphofructokinase, Afu4g00960), TCA (*idp1*; putative isocitrate dehydrogenase), HBP (*gfa1*; glutamine-fructose-6-phosphate transaminase, Afu6g06340), and PPP (glucose-6-phophate dehydrogenase (G6PDH), Afu3g08470) had increased transcript levels in *hrmA*^REV^ compared to AF293 (**Fig. 5D-G**). Importantly, transcripts of these rate limiting enzymes were reduced in *hrmA*^REV^Δ*uge3* compared to *hrmA*^REV^ (**Fig. 5D-G**). This suggests that flux through glycolysis, HBP, and PPP may be increased in *hrmA*^REV^ relative to AF293 and AF293Δ*uge3* strains, and this is consequently altered upon loss of *uge3* in *hrmA*^REV^.

Transcriptional differences observed in *hrmA*^REV^ such as the increased expression of key glycolytic, HBP, and TCA genes compared to AF293 shared overlap with metabolic alterations displayed by cancer cells in low oxygen environments (51). For example, increased transcript levels of genes in the HBP observed in *hrmA*^REV^ is a hallmark feature of cancer cells and used to generate nucleotide sugars that promote cell proliferation (**Fig. 5E**) (52, 53). Loss of *uge3* in *hrmA*^REV^ dramatically reduced the high transcript levels of the HBP rate limiting enzyme, Gfa1. We also observed significant transcript increases for genes in the PPP in *hrmA*^REV^ (**Fig. 5G**). In cancer, the PPP supports redox homeostasis through generation of NADPH that is required for cell anabolism and proliferation (54, 55). Further paralleling cancer metabolism, we observed increased transcripts encoding the TCA enzyme ATP-citrate lyase, *aclA*, in *hrmA*^REV^ compared to AF293 (**Fig. 5F**). Increased activity of ATP-citrate lyase in cancer supports proliferation through generation of acetyl-CoA for fatty acid synthesis (56, 57). Strikingly, transcripts of *aclA* are reduced upon loss of *uge3* in both AF293 and *hrmA*^REV^ suggesting Uge3 may impact TCA activity (**Fig. 5F**). Additionally, transcripts for several key enzymes involved in entry to and exit from glycolysis were increased in *hrmA*^REV^ compared to AF293 including: *hxkA* (hexokinase), *pfkA* (phosphofructokinase), and a putative pyruvate kinase (Afu6g07430), which strongly suggests *hrmA*^REV^ exhibits increased glycolytic flux (**Fig. 5D**). Importantly, transcripts for these genes were reduced upon loss of *uge3*, further suggesting that Uge3 plays a critical role in mediating this unique metabolic state (**Fig. 5D**). Taken together, these data provide transcriptional evidence that *uge3* plays a role in supporting a pseudohypoxic state in the H-MORPH strain, *hrmA*^REV^.

### Loss of the transmembrane glycosyltransferase, gtb3, results in GAG-deficiency but increases characteristics of H-MORPH

The finding that Uge3 plays a role in supporting the pseudohypoxic metabolic state exhibited by *hrmA*^REV^, led us to test the hypothesis that the growth and colony phenotypes observed in *hrmA*^REV^Δ*uge3* were due to *uge3’s* role in carbon metabolism versus the production of GAG. To test this hypothesis, we generated an *hrmA*^REV^Δ*gtb3* mutant and reconstituted strain, *hrmA*^REV^Δ*gtb3*^recon^. A *gtb3* null mutant has GAG-deficiency through loss of the transmembrane glycosyltransferase necessary for GAG production, but without disrupting *uge3* function (15). The *hrmA*^REV^Δ*gtb3* mutant and *hrmA*^REV^Δ*gtb3*^recon^ strains were confirmed via qRT-PCR and by crystal violet adherence assays (**Fig. S3A-B**). AF293Δ*gtb3* and AF293Δ*gtb3*^recon^ strains were generous gifts from the Sheppard Lab.

The loss of *gtb3* in the *hrmA*^REV^ background resulted in significant alterations to colony morphology, but did not impact colony growth in either strain parental strain background (**Fig. 6A-B).** Loss of *gtb3* in the AF293 background also did not impact colony morphology, meanwhile hrmA^REV^Δ*gtb3* exhibited increased characteristics of H-MORPH as defined by increased colony furrows and vegetative mycelia compared to its parental strain that was restored in the reconstituted strain (**Fig. 6C-D**). In submerged liquid culture, loss of *gtb3* did not impact total biomass produced in the AF293 background but resulted in a significant increase in total biomass produced in *hrmA*^REV^ (**Fig. 6E**). As expected, the Δ*gtb3* mutants in both AF293 and *hrmA*^REV^ had a complete lack of adherence that was restored with reconstitution of the wild-type allele (**Fig. 6F**). These data suggest that loss of *gtb3* also impacts colony morphology and growth of the H-MORPH strain *hrmA*^REV^ in contrast to the wild-type reference AF293, though in a different manner than loss of *uge3*.

**Figure 6.**
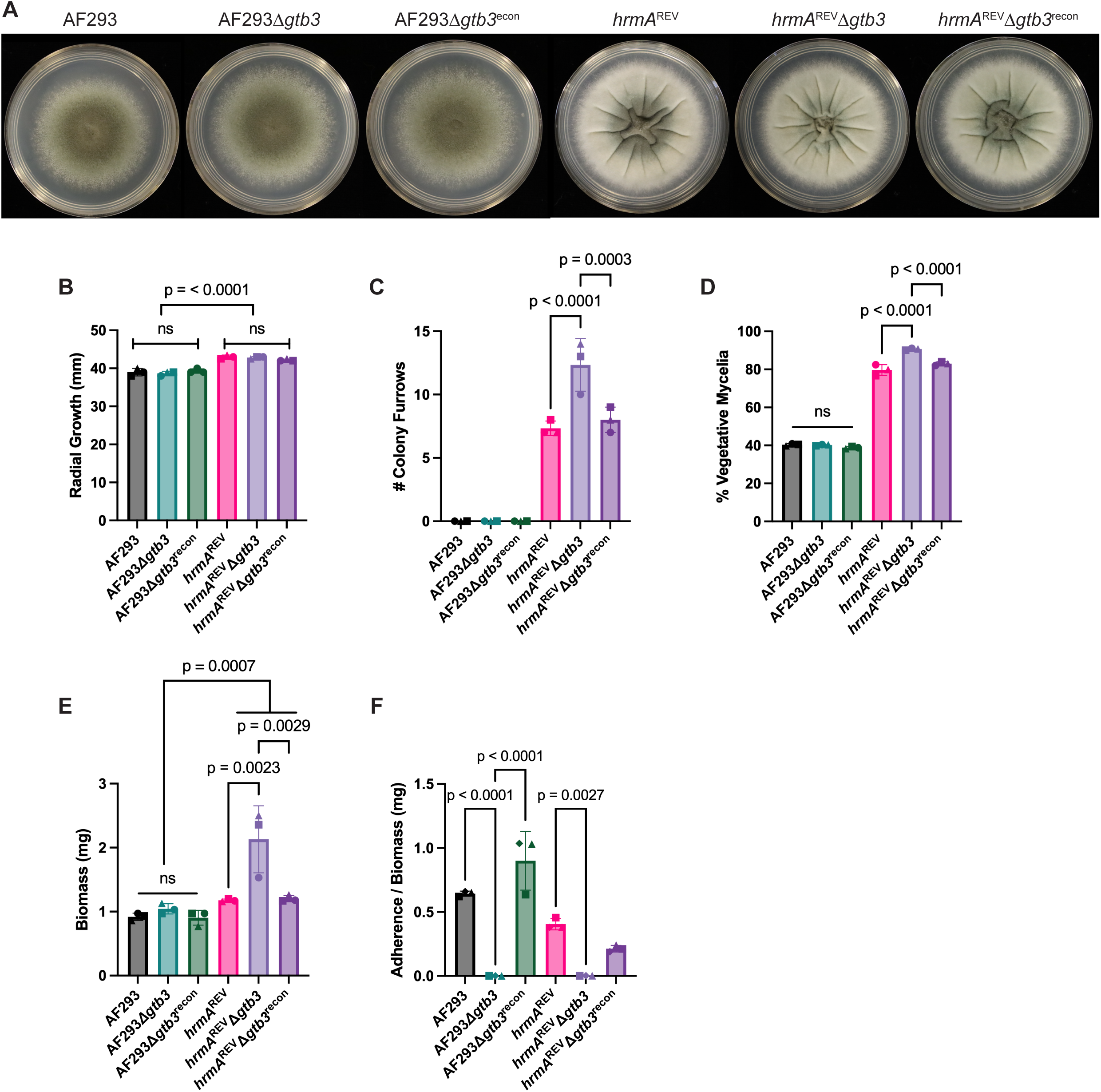
*hrmA*^REV^Δ*gtb3* displays increased characteristics of H-MORPH and increased biofilm biomass. **A-D)** Representative images of 72-hour colony biofilms utilized for quantifying impact to H-MORPH colony morphology via radial growth, number of colony furrows, and percent vegetative mycelia. Data depict three independent biological replicates. **E-F)** Paired biomass and crystal violet adherence data obtained from 24hr submerged biofilms (1x10^5^ spores/mL in LGMM). Data represents three independent biological replicates. **(B, E, F)** Statistical significance assessed via one-way ANOVA with Tukey’s multiple comparisons, **(C-D)** one-way ANOVA with Sidak’s multiple comparisons.

### HBP and PPP metabolism impact H-MORPH colony morphology

The finding that H-MORPH colony morphology in *hrmA*^REV^ was reduced upon loss of *uge3* and increased upon loss of *gtb3,* coupled with the increase in HBP and PPP associated transcripts in *hrmA*^REV^, suggested HBP generated nucleotide sugars and PPP reducing power play a critical role in modulating H-MORPH colony morphology and growth. To further address this hypothesis, we spot inoculated AF293, *hrmA*^REV^, and their respective GAG-deficient mutants (Δ*uge3* and Δ*gtb3*) on carbon-matched medias that enter central metabolism at specific points (**Fig. 7A**). We also included Sabouraud’s agar (SAB) as an undefined rich media as a control. Of note, this was the only media that was not carbon matched as it was prepared according to manufacturer specifications. We observed that on SAB all N-MORPH background strains exhibited the expected flat morphology, while all H-MORPH background strains including *hrmA*^REV^Δ*uge3* displayed furrowing (**Fig. 7B**). On all carbon matched medias, *hrmA*^REV^Δ*uge3* more closely phenocopied N-MORPH morphology than H-MORPH morphology. Interestingly, metabolic utilization of the oxidative phase of the PPP appears important for furrowing in H-MORPH, which is consistent with increased transcript levels of the PPP rate limiting enzyme, glucose-6-phosphate 1-dehydrogenase (Afu3g08470) in *hrmA*^REV^ (**Fig. 5G**). *hrmA*^REV^ background strains exhibited colony furrowing on carbon sources that can enter the oxidative phase of PPP, namely glucose and xylose (**Fig. 7B**). However, *hrmA*^REV^ background strains did not furrow when grown on fructose or glycerol, which bypass the oxidative phase of PPP or can enter in the non-oxidative phase of PPP, respectively (**Fig. 7B**). As the oxidative phase of the PPP is critical for NADPH generation and redox balancing, this suggests that *hrmA*^REV^ background colony furrowing is likely a physiological response to redox balancing via PPP (58).

**Figure 7.**
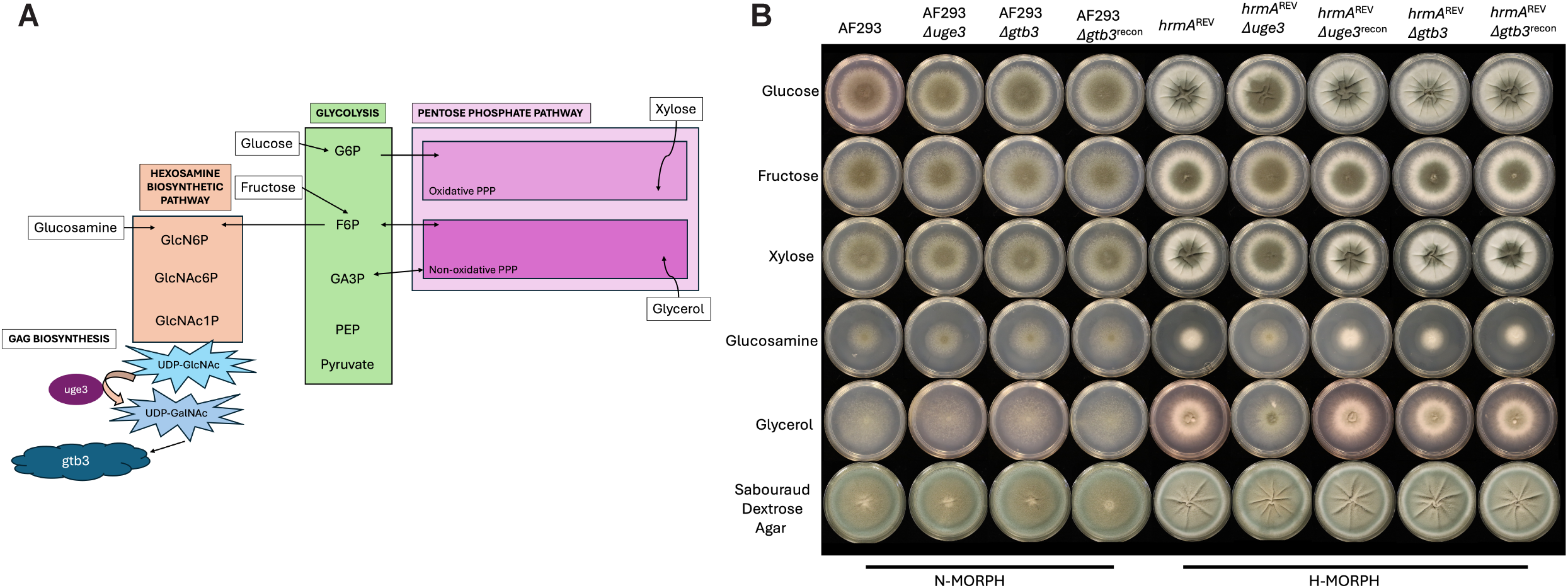
H-MORPH colony morphology is impacted by loss of *uge3* and relies on HBP and PPP metabolism. **A)** Model depicting where carbon sources tested in panel B would enter central carbon metabolic pathways, and suggested role of Uge3 downstream utilization of UDP-GlcNAc generated by HBP. **B)** 1x10^3^ spores in 2µL of 0.01% tween-80 from the indicated strains were plated onto 60mm petri dishes containing 10mL of carbon matched media, except SAB, which was utilized as an undefined, nutrient dense media. SAB was prepared to manufacturers specifications plus 1.5% agar as a solidifying agent. Plates were incubated for 72 hours at 37°C, 5% CO_2_, atmospheric O_2_, with humidity. Images are representative of three independent biological experiments.

Additionally, we observed less robust growth of all strains on glucosamine, but *hrmA*^REV^ background strains formed dense colonies while AF293 background strains formed “wispy” colonies, which is in concordance with increased transcriptional levels of HBP related genes in *hrmA*^REV^ compared to AF293 (**Fig. 5E**, **Fig. 7B**). Dense H-MORPH growth on glucosamine was disrupted by the loss of *uge3,* but not *gtb3,* suggesting *uge3* plays an essential role in mediating metabolism through the HBP in *hrmA*^REV^ (**Fig. 7B**). This coincides with a reduction of HBP related transcripts in *hrmA*^REV^Δ*uge3* compared to *hrmA*^REV^ (**Fig. 5E**). However, this was not particularly surprising as a main product of HBP is UDP-GlcNAc which is one of the substrates for Uge3 bifunctional epimerase activity (48, 59). Meanwhile, we did not observe morphological differences on glucosamine between AF293 background strains, which also is in alignment with low transcript levels of HBP related genes in AF293 and AF293Δ*uge3* (**Fig. 5E**, **Fig. 7B**). In combination with transcripts of the HBP rate-limiting enzyme, *gfa1* (Afu6g06340), being increased uniquely in *hrmA*^REV^, this data suggests *hrmA*^REV^ background strains utilize the HBP more than AF293 background strains, which is disrupted via the loss of *uge3*.

### Loss of uge3 impacts glucose uptake and carbon catabolite repression in H-MORPH

HBP and glycolysis are tightly linked to glucose metabolism and often function as nutritional stress sensors (60, 61). Furthermore, high glucose can increase the expression of *H. sapiens GFAT1* in cancer cells, suggesting a link between glycolytic flux and diversion of sugars into the HBP (62). A*s* we observed increased transcription in *hrmA*^REV^ of *pfkA*, encoding a rate-limiting enzyme in glycolysis, and *gfa1*, encoding a rate-limiting enzyme in HBP, we next asked if *hrmA*^REV^ background strains consumed more glucose than AF293 background strains, and if that was altered by the loss of *uge3*. To test this, we cultured our strains in LGMM in shake flask culture for 30 hours, then measured the glucose remaining in culture supernatant. We also collected biomass from the cultures as glucose utilization generates energy and provides carbon to support biomass production. From this experiment we observed that after 30 hours, AF293 background strains consumed very little glucose (**Fig. 8A**), and subsequently generated very little biomass (**Fig. 8B**). Of note AF293 mutant strains (Δ*uge3*, Δ*gtb3* and the reconstituted strain) all consumed significantly less glucose than AF293 by this timepoint. Meanwhile, *hrmA*^REV^ background strains consumed more glucose and produced significantly more biomass than AF293 by 30 hours (**Fig. 8A-B**). Interestingly, both loss of *uge3* and loss of *gtb3* in *hrmA*^REV^ resulted in significantly less glucose uptake and biomass production compared to the parental strain (**Fig. 8A-B**). However, *hrmA*^REV^Δ*gtb3* consumed more glucose and produced more biomass than *hrmA*^REV^Δ*uge3* (**Fig. 8A-B**). Taken together, these data suggest that *hrmA*^REV^ background strains consume more glucose relative to AF293 background strains that is utilized in anabolism to generate biomass and that *uge3* and *gtb3* are necessary for parental levels of glucose uptake and biomass production. These data highlight a central role in glucose metabolism for the GAG biosynthesis enzymes Uge3 and Gtb3.

**Figure 8.**
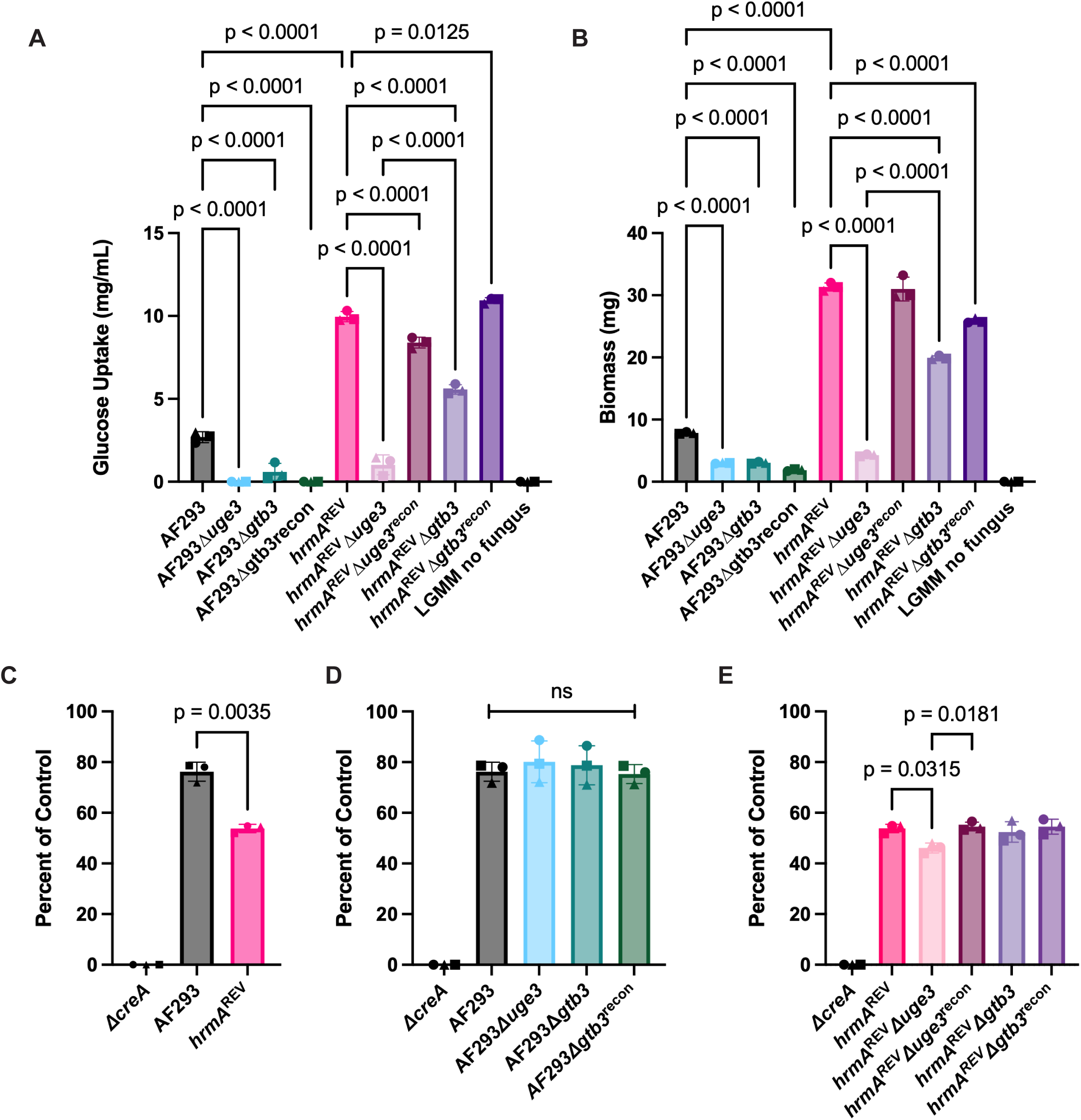
Loss of *uge3* impacts glucose uptake and carbon catabolite repression in H-MORPH strains. **A)** Glucose uptake from culture supernatant from 30 hour shaking cultures (1x10^6^ spores/mL in 10mL LGMM, 250rpm, 37°C, atmospheric O_2_, CO_2_). Glucose was measured from supernatant or LGMM no fungus control utilizing Glucose (GO) Assay Kit (Sigma Aldrich). Glucose values in mg/mL were extrapolated from a standard curve per manufacturer protocol, then glucose uptake was determined by subtracting the amount of glucose in the culture supernatant from the amount of glucose in the media only, no fungus control samples. Data represents three independent biological replicates. Statistical significance determined via one-way ANOVA with Sidak’s multiple comparisons. **B)** Fungal biomass produced from shake flask experiment described in panel A. Biomass from indicated cultures was collected in miracloth, washed with water, frozen at -80°C, then lyophilized prior to being weighed. Data represents three independent biological replicates. Statistical significance assessed via one way ANOVA with Sidak’s multiple comparisons. **C-E)** Glucose repression assay as previously described (64) 1x10^3^ spores in 2µL in 0.01% tween-80 from the indicated strains were plated onto 60mm petri dishes containing 10mL of either GMM or GMM + 0.1% allyl alcohol, then incubated for 48 hours (37°C, atmospheric CO_2_, atmospheric O_2_, with humidity). Sensitivity to allyl alcohol is a measurement of CCR de-repression, CEA10Δ*creA* was utilized as a positive control and exhibited 100% inhibition on GMM + 0.1% allyl alcohol. Data is representative of three independent biological experiments, statistical significance determined by one-way ANOVA with Tukey’s multiple comparisons.

Given the differences observed in glucose utilization and growth between AF293 and *hrmA*^REV^, and the transcript level differences in genes (e.g. *pfkA, alcB, alcC, adh1, facA, acuF, fbp1*) known to be directly or indirectly regulated by the carbon catabolite repressor, CreA (**Fig. 5 D,F,G**) (63), we hypothesized that carbon catabolite repression (CCR) is altered in *hrmA*^REV^ and is in part *uge3* dependent. We evaluated levels of CCR by plating the strains on either GMM or GMM with 0.1% allyl alcohol (AA), as previously described (64). Loss of *creA* in *A. fumigatus* results in CCR derepression resulting in full growth inhibition in the presence of AA, thus a Δ*creA* strain was used as a control (64). We observed that *hrmA*^REV^ exhibited increased sensitivity to AA compared to AF293, suggesting *hrmA*^REV^ is more CCR derepressed than AF293 (**Fig. 8C**). All AF293 background strains exhibited similar levels of CCR (**Fig. 8D**). Meanwhile, *hrmA*^REV^Δ*uge3* displayed a minor increase in sensitivity to AA compared to *hrmA*^REV^, while *hrmA*^REV^Δ*gtb3* displayed similar levels of CCR derepression to *hrmA*^REV^ (**Fig. 8E**). These data suggest that *hrmA*^REV^ displays increased CCR derepression relative to its AF293 parental strain and that loss of *uge3* but not *gtb3* further de-represses *hrmA*^REV^.

### GAG-deficient strains hrmA^REV^Δuge3 and hrmA^REV^Δgtb3 have divergent virulence phenotypes

In discovering *hrmA*^REV^Δ*uge3* occupies a metabolic state distinct from *hrmA*^REV^, it became unclear if the previously observed virulence defect upon loss of *uge3* was a consequence of the loss of GAG production or a result of metabolic dysregulation in central carbon metabolism. We hypothesized that if GAG produced by *hrmA*^REV^ contributes to virulence in this strain, then both GAG-deficient mutants, *hrmA*^REV^Δ*uge3* and *hrmA*^REV^Δ*gtb3,* would exhibit a reduction in virulence. To test this hypothesis, immune suppressed mice were challenged with either *hrmA*^REV^, *hrmA*^REV^Δ*gtb3*, *hrmA*^REV^Δ*gtb3*^recon^, or *hrmA*^REV^Δ*uge3*. While *hrmA*^REV^Δ*uge3* displayed a significant reduction in virulence as we had previously observed, *hrmA*^REV^Δ*gtb3* and *hrmA*^REV^Δ*gtb3*^recon^ did not exhibit a significant change in virulence (**Fig. 4A**, **Fig. 9A**). To better understand the dynamics of infection in *hrmA*^REV^Δ*gtb3*, we evaluated lung tissues at 3 days post challenge via histological examination (H&E and GMS). We had previously observed decreased fungal lesions present in the lung tissues of mice that received *hrmA*^REV^Δ*uge3* (**Fig. 4C**). In contrast, we observed larger foci of infection similar to *hrmA*^REV^ present in the lung tissues of mice challenged with either *hrmA*^REV^Δ*gtb3* or *hrmA*^REV^Δ*gtb3*^recon^ (**Fig. 9B**). These data suggest that *uge3* has a GAG-independent role that is necessary for virulence in the H-MORPH strain *hrmA*^REV^ and supports the hypothesis that the pseudohypoxic metabolic state of *hrmA*^REV^ is critical for its increased virulence in this murine model of IPA.

**Figure 9.**
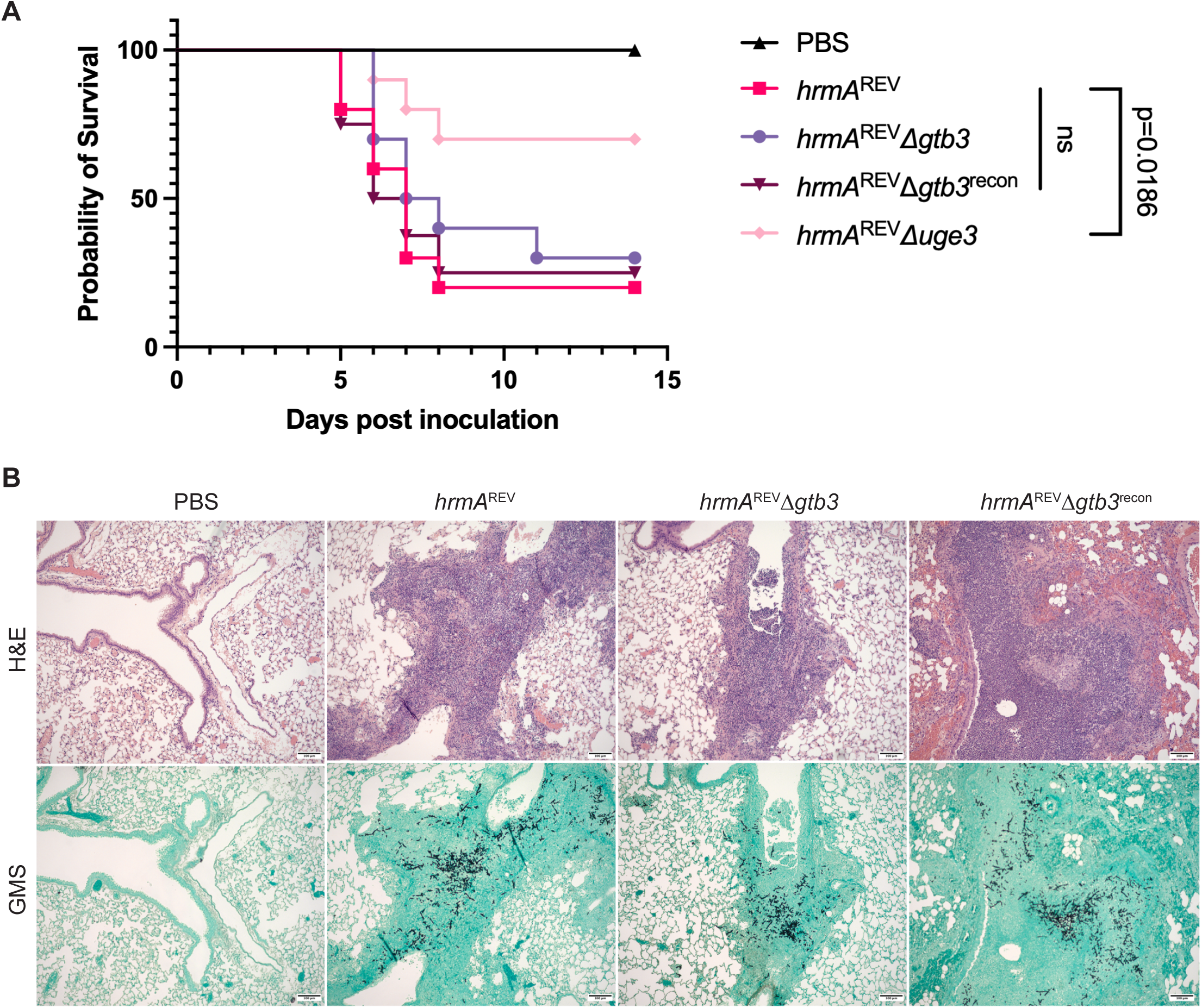
*hrmA*^REV^Δ*gtb3* and *hrmA*^REV^Δ*uge3* have divergent virulence phenotypes despite exhibiting GAG-deficiency. *hrmA*^REV^Δ*gtb3* and *hrmA*^REV^ Δ*uge3* are both GAG deficient and non-adherent, but only *hrmA*^REV^ Δ*uge3* displays a virulence defect. **A)** 6–8wk female CD-1 mice, were immunosuppressed with 40mg/kg triamcinolone acetonide (Bristol-Myers Squibb) 24 hours prior to intranasal inoculation with 1x10^5^ fungal spores of the indicated strains in 40µL PBS. Mice were monitored for signs of mortality and morbidity for 14 days. Data represents one biological experiment with n=10 mice/group. Statistical significance determined utilizing Kaplan-Meier survival analysis, Logrank (Mantel-Cox). **B)** Histopathology images (10X magnification; H&E and GMS) of corticosteroid immunosuppressed murine lungs 3 days post inoculation with 1x10^5^ fungal spores in 40µL PBS. Images are representative of one independent biological experiment with n=4 mice/group. Imaged on Zeiss Axioskop2 Plus utilizing SPOT (v5.2) imaging software, scale bars = 100µm.

## DISCUSSION

In this study, we describe a role for *uge3* in central carbon metabolism that is critical for the increased virulence displayed by the *A. fumigatus* H-MORPH strain *hrmA*^REV^ but not its parental N-MORPH strain AF293. Here we have provided evidence that increased mRNA levels of *hrmA* through the low oxygen evolved allele *hrmA*^D304G^ drives a carbon metabolic rewiring highlighted by transcriptional evidence of increased flux through glycolysis, and increased utilization of the HBP and PPP (**Fig. 5D-G**). This metabolic rewiring results in increased glucose consumption, partial CCR derepression, and relies on the oxidative phase of PPP to support increased growth and colony furrowing (**Fig. 8A, E**; **Fig. 7B**). The level of CCR derepression observed in *hrmA*^REV^ is further exacerbated in the Δ*uge3* mutant suggesting *uge3* plays a metabolic role in a strain-specific context (**Fig. 8E**). While it has been established that regulation of CCR through CreA is necessary for disease progression in the same murine model of IPA, further studies evaluating strains exhibiting a range of CCR are needed to discern how intermediate levels of CCR impact strain metabolic plasticity and contribute to virulence (64).

It was previously identified that the carbon catabolite repression regulator, CreA, positively regulates GAG production and as such Δ*creA* displays decreased adherence (65). Based on our current understanding of how CreA regulates central metabolism, it is not fully known how CreA positively regulates *uge3* but is suggested to be through modulating upstream regulators (63). Notably, loss of *uge3* in both AF293 and *hrmA*^REV^ resulted in decreased transcriptional levels of genes involved in the hexosamine biosynthetic pathway, except for *uap1* (**Fig. 5E**). Uap1 is a putative UDP-GlcNAc pyrophosphorylase which is the final step in UDP-GlcNAc synthesis but importantly can reversibly catalyze the first step of UDP-GlcNAc catabolism (59). UDP-GlcNAc is a critical metabolite involved in nutritional signaling; thus, modulating nucleotide sugar pools produced by the HBP could create a metabolic feedback loop to impact CCR, however this requires further investigation (66). Nonetheless, these data suggest a coordinated carbon utilization response is necessary to properly produce GAG, which is reduced in the H-MORPH *hrmA*^REV^.

A key question is why H-MORPH *hrmA*^REV^ is reliant on Uge3 activity for its robust growth and virulence. Across *Aspergillus* species, Uge3-like epimerases exhibit pleiotropic functions in metabolism and fungal physiology (44–47). Furthermore, the *H. sapiens* epimerase, GALE, is associated with galactose metabolism via the Leloir pathway. GALE deficiencies can lead to metabolic diseases such as galactosemia and lactose sensitivity in humans (67). *A. fumigatus* contains the genes involved in the Leloir pathway but *uge5* instead of *uge3* seems to be the major epimerase involved in galactose metabolism (19). In *H. sapiens*, GALE also exhibits a critical role in post-translational modifications and developmental regulation through protein glycosylation that when defective can dramatically impact human health (68, 69). Glycosylation as a post-translational modification is conserved in higher eukaryotes and is a critical function for growth, cell wall synthesis, and development in *A. fumigatus*, but it is not known if Uge3 is directly involved in glycosylation events (70–73).

Our transcriptional data and metabolic phenotyping of the H-MORPH strain *hrmA*^REV^ revealed patterns similar to low oxygen driven metabolic reprograming observed in cancer cells, a pseudohypoxic state. In cancer cells, cellular adaptation to low oxygen results in an increase in glycolytic metabolism to promote energy production and cellular proliferation (51). However, increased glycolytic flux, particularly if sustained over time, can result in oxidative stress and buildup of key glycolytic intermediates generating a cellular stress response (74). To help mitigate this stress, cancer cells can shunt glucose-6-phosphate into the pentose phosphate pathway and fructose-6-phosphate into the hexosamine biosynthetic pathway (53, 75, 76). In our experiments, we observed *hrmA*^REV^ consumed more glucose than AF293, which corresponded with increased biomass production (**Fig. 8A-B**). H-MORPH colony morphology could be diminished by providing carbon sources to the fungus that could not enter the oxidative phase of the pentose phosphate pathway, such as fructose and glycerol, suggesting the pentose phosphate pathway is involved in H-MORPH physiology. *hrmA*^REV^ displayed increased transcription of the rate-limiting enzymes involved in glycolysis, PPP, and HBP, which were reduced with the loss of *uge3* (**Fig. 5D-G**). Furthermore, H-MORPH colony morphology characteristics were reduced through loss of *uge3*, which utilizes UDP-GlcNAc generated via the hexosamine biosynthetic pathway, also suggesting a role for HBP in H-MORPH physiology (**Fig. 3A-D**). Interestingly, we observed an increase in H-MORPH characteristics in the absence of *gtb3* suggesting that enzymatic activity of Uge3 is involved in H-MORPH physiology, distinct from but related to its known role in GAG synthesis (**Fig. 6A-D**). Consequently, one potential model posits that flux through HBP is increased in *hrmA*^REV^, and Uge3 epimerase activity is needed to manage the rapidly produced UDP-GlcNAc.

In *B. subtilis*, loss of the UDP-galactose-4-epimerase, GalE, resulted in accumulation of UDP-Gal causing a toxic effect (77). It is possible that loss of *uge3* in *hrmA*^REV^ causes accumulation of UDP-GlcNAc generated from HBP, causing some form of cellular toxicity and/or metabolic feedback that limits metabolism and reduces growth. For example, loss of the carbon catabolite repressor CreA in *A. fumigatus* reduces growth on all tested carbon sources to date, which is hypothesized to be due to a futile metabolic cycle that ultimately depletes anaplerotic reactions needed for growth (64). In alignment with this possibility, we observed that loss of *uge3* in *hrmA*^REV^ resulted in a significant growth defect in colony biofilms, a reduction in CCR, and these phenotypes were not observed upon loss of *gtb3* which retains Uge3 activity (**Fig. 3A-B**; **Fig. 6A-B**). Additionally, as *uge3* and *gtb3* null *hrmA*^REV^ strains exhibited different virulence phenotypes this suggests the metabolism role of Uge3 is important for virulence in a strain-specific context. However, from our studies it is not fully understood if Uge3 activity mitigates cellular toxicity of the buildup of UDP-GlcNAc, or promotes growth by epimerizing UDP-GlcNAc for utilization in an unidentified process other than GAG production.

Our data suggest *hrmA*^REV^ background strains more rapidly accumulate biomass than AF293 background strains, and according to our transcriptional data, may adopt similar cancer-like patterns of metabolic behavior to facilitate this biomass production (**Fig. 8B**; **Fig. 5D-G**) (76). However, unlike in comparisons between cancer cells and non-cancer cells, our glucose consumption per biomass produced seems to be comparable between our H-MORPH and our N-MORPH strains (**Fig. 8A-B**). Supporting that, altered metabolic flux and the resulting energy deficiency that arises from prioritizing biomass production rather than ATP production is supposedly a major driver of the increased glucose uptake observed by cancer cells (76, 78–80). It remains an open question how *hrmA*^REV^ background strains might be maintaining that balance, should this framework be accepted.

Perhaps the answer lies outside of the cytoplasm. We have previously observed H-MORPH strains possess a thinner cell wall, a structure primarily composed of beta-glucan, a glucose derived molecule, and chitin, which is composed of N-acetyl glucosamine (11). Additionally, we have shown *hrmA*^REV^ exhibits a significant reduction in total extracellularly secreted sugars, particularly a reduction in GAG monosaccharides (Gal and GalNAc) (**Fig. 2A**). Rather than increasing per-cell import, *hrmA*^REV^ appears to be decreasing sugar export. This reprioritization of sugars away from the cell wall and GAG production may serve to support the increased growth rate observed in these strains (**Fig. 2A**).

Furthermore, loosening of CCR may be an adaptation in line with *A. fumigatus’* environmental niche as a saprophyte, and facilitate expansion of acceptable carbon sources to facilitate increased vegetative growth (**Fig. 8C-E**). It is also fascinating to consider that the oxidative PPP seems integral to the development of the characteristic furrows in the H-MORPH colony (**Fig. 7B**). The PPP is a major mechanism of NADPH regeneration, a metabolite critical for the production of fatty acids, sterols, and nitrate reduction (54, 55, 58). Future experiments testing various aspects of this framework will provide insights into the causes and consequences underlying the observations made here.

In summation, our data supports that the low oxygen evolved allele *hrmA*^D304G^ (*hrmA*^REV^) is sufficient to induce a pseudohypoxic metabolic reprogramming similar to what is observed in cancer. As this allele and strain were selected for under a high glucose, high nitrate, low oxygen environment, these data are consistent with the conditions the strain arose in. Notably, as all experiments in this study were conducted at atmospheric oxygen tensions (∼21%), *hrmA*^D304G^ can induce this pseudohypoxic metabolic program independent of oxygen, similar to what has been described in cancer cells (81, 82). Thus, we propose that *hrmA* mediated H-MORPH drives a pseudohypoxic response, increasing glycolysis, and shunting sugars to parallel metabolic pathways HBP, and PPP. The pseudohypoxic response alters colony morphology via the PPP and promotes proliferation and virulence in part through the activity of Uge3 (**Fig. 10**).

**Figure 10.**
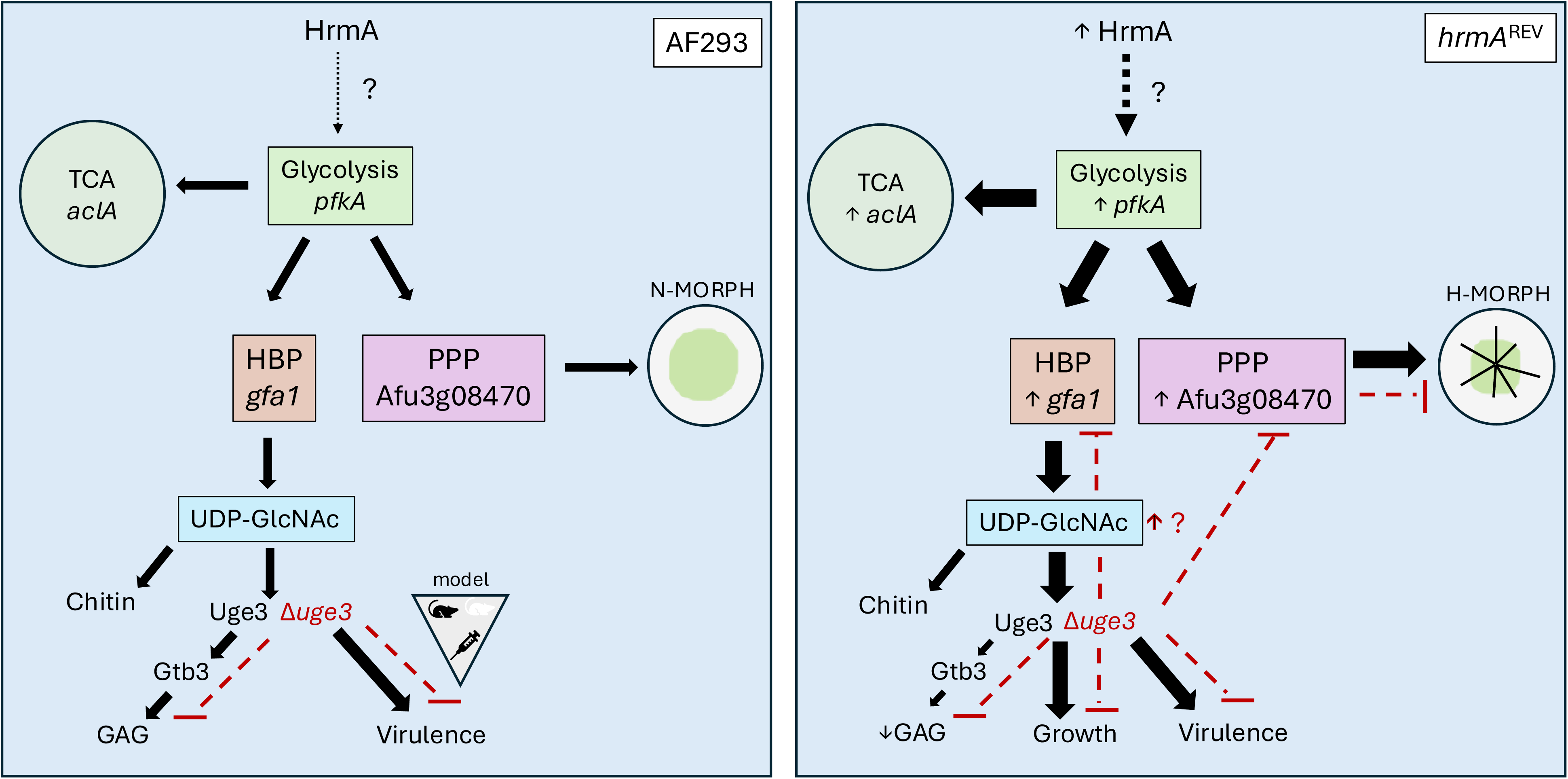
Model depicting the relationship between the pseudohypoxia metabolic state of H-MORPH *hrmA*^REV^ and Uge3 in mediating colony morphology, growth, and virulence. The nature of the interaction between increased *hrmA* expression and the impact to glycolysis is unknown. Increased *hrmA* expression also results in increased TCA (*aclA*), PPP (Afu3g08470, putative glucose-6-phosphate dehydrogenase (G6PD), and HBP (*gfa1*). PPP activity promotes colony furrowing (H-MORPH). HBP activity generates UDP-GlcNAc that supports synthesis of chitin for the cell wall and promotes the growth and virulence of *hrmA*^REV^ but is diverted from use in GAG production. Loss of *uge3* in *hrmA*^REV^ results in a growth defect, reduced transcripts of *gfa1,* reduced transcripts of the putative G6PD that coincides with reduced H-MORPH, and reduced virulence. In AF293, *hrmA* is lowly transcribed in lab conditions, and as such TCA, HBP, and PPP activity are at a baseline of activity. This amount of PPP activity results in a flat (N-MORPH) colony morphology. UDP-GlcNAc produced by HBP activity supports synthesis of chitin for the cell wall and is utilized for GAG production. Presence or absence of *uge3* in AF293 does not significantly alter growth or virulence in a triamcinolone immune suppressed murine model of IPA, however model of IPA can influence virulence phenotype.

Further investigation into how sugars epimerized by Uge3 contribute to virulence outside of the scope of GAG production, and how HrmA induces a pseudohypoxic cellular state could yield new insights for IPA therapies and also be valuable for fungal biotechnology applications.

## METHODS AND MATERIALS

### Strains and growth conditions

Mutant strains were made in the *Aspergillus fumigatus* AF293 laboratory reference strain. Strains were stored as conidia in 25% glycerol at −80°C and maintained on 1% glucose minimal medium [GMM; 6 g/liter NaNO_3_, 0.52 g/liter KCl, 0.52 g/liter MgSO_4_·7H_2_O, 1.52 g/liter KH_2_PO_4_ monobasic, 2.2 mg/liter ZnSO_4_·7H_2_O, 1.1 mg/liter H_3_BO_3_, 0.5 mg/liter MnCl_2_·4H_2_O, 0.5 mg/liter FeSO_4_·7H_2_O, 0.16 mg/liter CoCl_2_·5H_2_O, 0.16 mg/liter CuSO_4_·5H_2_O, 0.11 mg/liter (NH_4_)6Mo_7_O_24_·4H_2_O, 5 mg/liter Na_4_EDTA, 10g/liter glucose, pH 6.5]. Solid medium was prepared by addition of 1.5% agar prior to autoclaving, while experiments that called for liquid media, agar was not added. For all experiments, *A. fumigatus* was grown on solid GMM at 37°C, 5% CO_2_, with humidity for 3 days to produce conidia. Conidia were collected using 0.01% Tween 80 and filtered through miracloth, counted using a hemacytometer, and diluted in either 0.01% Tween-80 or medium to the final concentration used in each assay.

All experiments were performed with GMM unless explicitly stated otherwise. For experiments where carbon sources were changed, the quantity of carbon was matched to the total carbon in GMM (e.g. 2% glycerol (20g/L), 1.2% xylose (12g/L), 1% fructose (10g/L), 1% glucosamine (10g/L)), but nitrate was still utilized as the nitrogen source. SAB (Difco; ref# 23820) was prepared according to the manufacturer’s instructions and 1.5% agar was used as the solidifying agent, notably this media was not carbon matched.

### Strain generation

All strains utilized in this study are listed in **Table S2**. The laboratory reference strain AF293 and the isogenic H-MORPH strain *hrmA*^REV^ were used as wildtypes throughout these studies (11, 83). AF293Δ*uge3*, AF293Δ*gtb3*, and AF293Δ*gtb3*^recon^ strains were a generous gift of the Sheppard Lab (McGill University, Montreal, QC). For generation of *hrmA*^REV^Δ*uge3* the open reading frame (Afu3g07910) was replaced with the *hphB* selection marker. The replacement construct was generated using overlap PCR to fuse ∼1 kb upstream and ∼1 kb downstream of the open reading frame of *uge3* to the *hphB* marker. The resulting construct was transformed into protoplasts from *hrmA*^REV^, and transformants were selected on SMM (GMM + 1.2M sorbitol) with 175 μg/mL hygromycin B (VWR). The reconstituted strain was generated by amplifying from ∼1 kb upstream of the start codon to ∼1 kb downstream of the stop codon, then ligating that construct into a plasmid containing the *bleoR* resistance marker. The generated plasmid was ectopically transformed into protoplasts from *hrmA*^REV^Δ*uge3*, and transformants were selected on SMM with 100µg/mL phleomycin (Invivogen). Strains were confirmed via PCR, qRT-PCR, and adherence assays. Transformation into protoplasts was conducted as previously reported (84). For generation of *hrmA*^REV^Δ*gtb3*, CRISPR-Cas9 mediated transformation was utilized to replace the *gtb3* (Afu3g07860) open reading frame of with the *hphB* selection marker (85). Transformation into protoplasts and selection of transformants was conducted as described above. For the reconstituted strain ∼1.5kb upstream of the start codon and ∼400bp downstream of the stop codon were amplified using primers with microhomology to a plasmid backbone containing a *bleoR* selection marker, then the construct was ligated to the plasmid backbone using NEBuilder Hi-Fi DNA assembly kit (NEB). The repair construct was amplified from the resulting plasmid using primers with microhomology to the *aft4* safe haven site, then transformation was carried out using CRISPR-Cas9 (85, 86). Transformants were selected on SMM containing 100µg/mL phleomycin (Invivogen). Successful mutants and reconstituted strains were confirmed via PCR, qRT-PCR, Southern blot, and adherence assays.

### In vitro growth and adherence assays

For radial growth assays 1000 fungal spores in 2µL of 0.01% Tween-80 onto plates and incubated at 37°C, 5% CO_2_, with humidity for 72 hours. Macroscopy morphology quantifications were conducted as described previously (11). Biomass assays were conducted by inoculating 3mL of 1x10^5^ spores/mL in liquid glucose minimal media (LGMM) in technical triplicate into wells of a 6-well plate. Cultures were statically incubated (37°C, 5% CO_2_, with humidity) for 24 hours prior to collection. Collected biomass was washed twice with water, frozen at -80°C, then lyophilized and weighed. Crystal violet adherence assays were conducted as described previously (84). Carbon catabolite repression assays were also conducted as described previously (64).

For quantification of glucose uptake experiments, for each strain 1x10^6^ spores/mL in 10mL of LGMM was incubated for 30 hours, shaking at 250rpm, incubated at 37°C (atmospheric O_2_/CO_2_). After incubation, fungal biomass and culture supernatants were separated by filtration through miracloth. Fungal biomass was washed with water, frozen at -80°C, then lyophilized prior to weighing. To quantify glucose uptake, we used the Glucose (GO) Assay Kit (Sigma) using the following modifications. 1mL of homogenized fungal culture supernatant was incubated for 10 minutes in a 65°C waterbath to thermally inactivate any native glucose oxidase, as the glucose oxidase utilized in the Glucose (GO) Assay Kit (Sigma) originates from *Aspergillus niger* (87).

Incubated culture supernatants were then diluted 1:100 in molecular grade biology water to fall in the range of detection for the kit. We followed the manufacturer assay protocol with the volume modifications for a 96-well format.

### Derivatization and gas chromatography mass spectrometry (GC-MS) to determine ECM composition

To obtain secreted extracellular oligosaccharides, whole culture supernatants from shake flask cultures (100mL of 1x10^6^ conidia/mL, 200rpm, 37°C for 42 hours) were dialyzed against water in a 3.5kDa MWCO bag (Spectrum Labs, #132725) for three days with water being changed 3x/day. Resulting dialyzed culture supernatants were lyophilized prior to analysis via GC-MS. Lyophilized samples and internal standards, and calibrations sets were prepared following an O-Methyl Trimethylsilyl monosaccharide derivative composition analysis protocol as described (38). The resulting TMS methyl glycosides were dried, resuspended in 1 ml of cyclohexane, and injected into the GC-MS system (Agilent Technologies GC7890B and MS Ensemble 5977B) equipped with a CP-Sil5-CB capillary column (Agilent Technologies CP7441). Elution and acquisition were performed as described. Identification and quantification of each monosaccharide were carried out using standards and response factors determined for each monosaccharide.

### RNA Extraction and qRT-PCR

For RNA-sequencing and qRT-PCR, mycelia from 24 hour submerged liquid cultures (1x10^5^ conidia/mL in liquid glucose minimal media) were collected in TRIsure (Sigma) and kept on ice, then and bead beat for 1 minute with 2.3 mm beads. Homogenate was brought to a total volume of 1 mL TRIsure and RNA was extracted as previously described (64). For qRT-PCR and RNA-sequencing, 5µg of RNA was DNAse treated with Ambion Turbo DNAse (Life Technologies) according to the manufacturer’s instruction. For qRT-PCR DNase treated-RNA was processed as previously described (64). mRNA levels were normalized to *actA* and *tefA* for all qRT-PCR analyses. qRT-PCR data was collected on a CFX Connect Real-Time PCR Detection System (Bio-Rad) with CFX Maestro Software (Bio-Rad).

### RNA-Sequencing: Library Preparation and Sample Sequencing

For RNA-Seq, RNA was quantified by Qubit (Thermo Fisher Scientific). Library preparation and sample sequencing were conducted using SeqCoast Genomics (Portsmouth, NH, USA). Briefly, RNA samples were prepared for sequencing using the Illumina Stranded mRNA Prep (#20040534) with Illumina Unique Dual Indexes.

Sequencing was performed on the Illumina NextSeq2000 platform using a 300-cycle XLEAP-SBS flow cell kit to produce 2x150bp paired reads. 1-2% PhiX control was spiked into the run to support optimal base calling. Read demultiplexing, read trimming, and run analytics were performed using DRAGEN v4.2.7, an on-board analysis software on the NextSeq2000. Files were generated as fastq.gz files for downstream analyses.

### RNA-Sequencing: data processing and differential expression analysis

RNA-Seq data processing was conducted in-house. Raw sequencing reads were quality assessed using FastQC v0.11.8 and summarized with MultiQC v1.10.1 (88, 89). Adapter sequences and low-quality bases were trimmed using Cutadapt v2.4 (--nextseq-trim=20, -q 20, -m 1) with poly-A tail removal (90). Trimmed reads were re-evaluated with FastQC and MultiQC prior to alignment. Reads were aligned to the AF293 reference genome (FungiDB v52) using STAR v2.7.2b with coordinate-sorted BAM output (91–93). Alignment quality was assessed with MultiQC. For quantification, BAM files were name-sorted using SAMtools v1.9 and gene-level read counts were generated with HTSeq-count v0.11.2 (94, 95). Per-sample count files were combined into a single count matrix for downstream analysis.

For the exploratory analysis, DESeq2 v1.50.2 was used for generating variance stabilizing transformed (VST) counts following removal of lowly expressed genes (96). Sample relationships were assessed by hierarchical clustering of Euclidean distances on VST-normalized counts. Differential expression analysis used the limma package v4.8.2 using limma-voom methodology (97). Counts were filtered using EdgeR v4.8.2 filterByExpr function and normalized using the trimmed mean of M-values (TMM) method (98). Empirical Bayes moderation was applied and pairwise contrasts were tested between strains. Heatmaps were generated using the ComplexHeatmap package (99, 100). Kyoto Encyclopedia of Genes and Genomes (KEGG) analysis used the KEGGREST package v1.50.0 and pathview v1.50.0 (101, 102).

### Murine Survival Experiments

All animal experiments were performed following experimental methodology as described (11). In brief, 6–8-week-old female CD-1 mice (Charles River Laboratory) weighing between 24-28g were immune-suppressed via subcutaneous injection with 40mg/kg triamcinolone acetonide (Kenalog-10; Bristol-Myer Squibb) 24 hours prior to fungal challenge. For fungal challenge, mice were lightly anesthetized with isoflurane, then intranasally instilled with 1x10^5^ fungal conidia in 40µL of PBS, and mock mice received 40µL PBS. Mice were maintained under a 12:12 light dark cycle, with food and water *ad libitum*, then monitored for 14 days post challenge for signs of morbidity and mortality. Kaplan-Meier curves were generated and Log-rank Mantel-Cox tests and Gehan-Breslow-Wilcoxon tests performed.

For Dectin-1 murine experiments, wildtype C57BL/6J (stock #000664; Jackson Laboratory) and *Clec7a*^-/-^ mice (stock #012337, Jackson Laboratory) were backcrossed and wildtype and knockout breeders were established to maintain the colony in house. Genotypes were confirmed utilizing provided Jackson Labs genotyping protocol for stock #012337. For all subsequent experimentation, 8-12-week-old, age, sex, and littermate matched mice were utilized, then mice were immune-suppressed and challenged with fungal conidia as described above.

### Histology and Fungal Burden via qPCR

For histopathological analyses mice were immune-suppressed and challenged with fungus as described above. Lungs were harvested on 3 days post-inoculation and were prepared for Gömöri methenamine silver (GMS) and hematoxylin and eosin (H&E) staining or fungal burden quantification as described (64). Images of tissue-matched samples were taken using Zeiss Axioskop2 Plus utilizing SPOT (v5.2) imaging software. Scale bars were added using ImageJ2 (v2.16.0/1.54p) (103).

For fungal burden via qPCR, lungs were extracted at 3 days post inoculation and lung tissue was flash frozen, then freeze-dried and homogenized with 2.3mm zirconia/silica beads on a Mini-BeadBeater (BioSpec Products, Inc.). Total DNA was extracted using the Plant and Fungal DNA kit (Omega Bio-tek) and RNAse treated DNA was used for quantitative PCR, utilizing primers to amplify 18S rRNA as previously described (104).

### Statistical Analyses

All statistical analyses were conducted in GraphPad Prism 11, error bars depict standard deviation. Non-parametric analyses were utilized for all murine experiments.

## Data Availability Statement

RNA sequencing data that support the findings of this study have been deposited under NCBI Gene Expression Omnibus with the identifier GSE334031under the BioProject PRJNA1473623. All strains and any other data used to support these findings will be made available upon reasonable request to the corresponding author.

## AUTHOR CONTRIBUTIONS

NEK generated strains utilized in the studies, designed and performed experiments, analyzed and interpreted the data, and wrote the manuscript. CP conducted RNA-Seq analysis, aided in data interpretation, and generated figure panels presented in Figure 5. JTJ and KGQ contributed to murine studies. CHK and KWL aided in strain generation. FLM conducted GAG compositional analyses and helped interpret the data. AJ contributed to the conceptual framework and helped write the manuscript. RAC provided funding, contributed to experimental design, data interpretation, and helped write the manuscript.

## ACKNOWLEDGEMENTS

We would like to thank and acknowledge the Sheppard lab for the gifting of strains AF293Δ*uge3*, AF293Δ*gtb3*, and AF293Δ*gtb3*^recon^. This work was funded by NIH/NIAID grant R01AI181215 (RAC). NEK was also supported by NIH/NHBLI 5T32HL134598 grant (O’Toole, George) and by NIH/NIAID 1F31AI88661 (Kordana, Nicole). Core facility support provided by NIH grant P20-GM113132 to the Dartmouth BioMT COBRE. Shared Resources Center pathology services support provided through NCI Cancer Center Support Grant 5P30CA023108 (RRID: SCR_023479). Molecular Interaction and Imaging Support Core support provided through NIH IDeA award for Dartmouth BioMT P20-GM113132. Polysaccharide analyses were supported through GlycoNet Integrated Services financed by the Canadian Fund for Innovation, major sciences initiative program (Grant 42481).

## SUPPLEMENTARY FIGURE LEGENDS

**Supplementary Figure 1. Confirmation of *hrmA*^REV^Δ*uge3* and reconstituted strain. A)** qRT-PCR data of *uge3*. RNA was extracted from 24hr submerged culture biofilms (1x10^5^ spores/mL in LGMM) incubated in 5% CO_2_, atmospheric O_2_, 37°C, with humidity. Cq values were normalized to *actA* and *tefA*, then *uge3* expression in the mutant strains was normalized to *uge3* expression in the wildtype, *hrmA*^REV^. Data represents one biological replicate averaging three technical replicates. Reconstituted strain was selected by exhibiting best restoration of expression and adherence of all selected transformants. **B)** Crystal violet adherence data from 24hr static liquid submerged biofilms (1x10^5^ spores/mL in LGMM). Data represents three independent biological replicates. Significance was assessed via one-way ANOVA with Tukey’s multiple comparisons.

**Supplementary Figure 2. RNA-Seq quality assurance indicates adequate sample clustering. A)** Sample relationships visualized by hierarchical clustering of Euclidean distances on VST-normalized counts indicates overall good RNA-Sequencing quality with few samples exhibiting variation.

**Supplementary Figure 3. Confirmation of *hrmA*^REV^Δ*gtb3* and reconstituted strain. A)** qRT-PCR data of *gtb3*. RNA was extracted from 24hr submerged culture biofilms (1x10^5^ spores/mL in LGMM) incubated in 5% CO_2_, atmospheric O_2_, 37°C, with humidity. Cq values were normalized to *actA* and *tefA*, then *gtb3* expression in the mutant strains was normalized to *gtb3* expression in the wildtype, *hrmA*^REV^. Data represents one biological replicate averaging three technical replicates. Reconstituted strain was selected by exhibiting best restoration of expression and adherence of all selected transformants. **B)** Crystal violet adherence data from 24hr static liquid submerged biofilms (1x10^5^ spores/mL in LGMM). Data represents three independent biological replicates. Significance was assessed via one-way ANOVA with Tukey’s multiple comparisons.

